# Desiccation resistance differences in *Drosophila* species can be largely explained by variations in cuticular hydrocarbons

**DOI:** 10.1101/2022.06.24.497513

**Authors:** Zinan Wang, Joseph P. Receveur, Jian Pu, Haosu Cong, Cole Richards, Muxuan Liang, Henry Chung

## Abstract

Maintaining water balance is a universal challenge for organisms living in terrestrial environments, especially for insects, which have essential roles in our ecosystem. Although the high surface area to volume ratio in insects makes them vulnerable to water loss, insects have evolved different levels of desiccation resistance to adapt to diverse environments. To withstand desiccation, insects use a lipid layer called cuticular hydrocarbons (CHCs) to reduce water evaporation from the body surface. It has long been hypothesized that the waterproofing capability of this CHC layer, which can confer different levels of desiccation resistance, depends on its chemical composition. However, it is unknown which CHC components are important contributors to desiccation resistance and how these components can determine differences in desiccation resistance. In this study, we used machine learning algorithms, correlation analyses, and synthetic CHCs to investigate how different CHC components affect desiccation resistance in 50 *Drosophila* and related species. We showed that desiccation resistance differences across these species can be largely explained by variation in cuticular hydrocarbons. In particular, length variation in a subset of CHCs, the methyl-branched CHCs (mbCHCs), is a key determinant of desiccation resistance. We also showed a significant correlation between the evolution of longer mbCHCs and higher desiccation resistance. Given the ubiquitous presence of mbCHCs in insects, the evolution of mbCHCs may be a general mechanism of how insects evolve desiccation resistance and adapt to diverse and changing environments.

**Significance:** As our planet is becoming more arid due to global warming, preventing dehydration is key to the survival of insects, an essential part of our ecosystem. However, factors that determine how insects may evolve resistance to desiccation are relatively unknown. Using *Drosophila* species from diverse habitats, we showed that variations in the composition of cuticular hydrocarbons (CHCs), a hydrophobic layer found on insects to prevent evaporative water loss, can largely explain desiccation resistance differences. In addition, the evolution of longer methyl-branched CHCs (mbCHCs), underlies the evolution of higher desiccation resistance in this genus. As mbCHCs are ubiquitously present in most insects, we suggest that evolutionary changes in mbCHCs may be a general determinant of desiccation resistance across insect species.

## Introduction

Maintaining water balance is a universal challenge for organisms living in terrestrial environments where water constantly evaporates from the body surface (Hadley, 1994). Organisms such as insects have a high surface area to volume ratio due to their small sizes, rendering them vulnerable to water loss and desiccation (Gibbs, 2002b; Gibbs and Rajpurohit, 2010; Kühsel et al., 2017). Studies using closely related insect species collected worldwide showed that phylogenetically related insect species can have very different levels of desiccation resistance and occupy very distinct habitats, while insects that are phylogenetically distant but dwelling in similar habitats could have similar levels of desiccation resistance (Kellermann et al., 2012; Li et al., 2022; Menzel et al., 2017; Rane et al., 2019). This suggest that extant species have evolved different levels of desiccation resistance to survive in their different habitats. However, as climate change accelerates the expansion of dryland (Huang et al., 2016) and changes aridity in many areas across the globe (Sherwood and Fu, 2014; Shi et al., 2021), it is less understood how insect species, which are integral parts of our ecosystems, can evolve higher levels of desiccation resistance to adapt to the more arid environments.

To determine how insects can adapt to desiccation stress, understanding how insects conserve water and prevent desiccation is crucial. In insects, cuticular water loss is the leading cause of desiccation (Chown et al., 2011; Gibbs and Matzkin, 2001; Wang et al., 2021). In an extreme example, cuticular water transpiration has been found to account for 97% of increased water loss in queens of the harvester ant *Pogonomyrmex barbatus* (Johnson and Gibbs, 2004). To conserve water and prevent desiccation, a general mechanism in insects is the use of a lipid layer on the epicuticle, named cuticular hydrocarbons (CHCs) (Gibbs et al., 2003; Gibbs and Matzkin, 2001). This CHC layer, which can contain more than 100 different compounds on the same individual, provides a hydrophobic barrier against evaporative water loss through the cuticle (Blomquist and Ginzel, 2021). This was first demonstrated when the physical or chemical removal of this layer using abrasive dust and various solvents resulted increased water loss in various insect species in experiments published almost eight decades ago (Wigglesworth, 1945). In recent years, after the identification of a cytochrome P450 decarbonylase responsible for the synthesis of insect CHCs (Qiu et al., 2012), genetic manipulations of this gene in the fruitfly *Drosophila melanogaster* as well as in aphids and cockroaches, led to almost complete loss of CHCs and significant decrease in desiccation resistance (Chen et al., 2016; Chen et al., 2019; Qiu *et al*., 2012).

Variations in the composition of this CHC layer has been suggested to contribute to intraspecific variation in desiccation resistance in *D. melanogaster* (Rouault et al., 2004), the Mediterranean dung beetle *Onthophagus taurus* (Leeson et al., 2020), and the Argentine ant *Linepithema humile* (Buellesbach et al., 2018). This is supported by studies manipulating CHC composition either chemically or genetically, in different *Drosophila* species, producing different levels of desiccation resistance (Chiang et al., 2016; Chung et al., 2014; Ferveur et al., 2018; Koto et al., 2019; Savage et al., 2021). Together, these studies suggest that different components in the CHC layer can influence desiccation resistance.

The varying ability of these CHC components to prevent desiccation depends on its chemical structure, which in turn determines its melting temperatures (Gibbs and Pomonis, 1995; Gibbs, 1998; 2002a). The melting temperature of the hydrocarbon is positively correlated with its water-proofing properties and contribution to desiccation resistance (Blomquist and Ginzel, 2021; Gibbs and Rajpurohit, 2010). In *Drosophila* and most other insects, CHCs range in lengths between approximately 21 to 50 carbons, and consist of linear alkanes (*n*-alkanes), alkenes (monoenes and dienes), and methyl-branched alkanes (Blomquist and Ginzel, 2021). Among them, *n*-alkanes have the highest melting temperature, followed by methyl-branched alkanes (mbCHCs), monoenes, and dienes. Increase in CHC chain length can also increase melting temperature and potentially leads to higher desiccation resistance (Gibbs and Pomonis, 1995). This is consistent with laboratory selection experiments in *D. melanogaster* selecting for increased desiccation resistance resulted in longer carbon-chain CHCs in the desiccation selected flies than the control flies (Gibbs et al., 1997; Kwan and Rundle, 2010). Longer CHC chain length is also correlated with climatic factors such as higher temperature and lower precipitation (Jezovit et al., 2017; Rouault *et al*., 2004). These climatic factors are associated with desiccation resistance in *Drosophila* species (Hoffmann, 2010; Hoffmann and Weeks, 2007; Kellermann et al., 2018).

While the studies mentioned above showed strong evidence that CHCs and variations in CHC compositions are associated with desiccation resistance in insects, no study has investigated the extend that CHC variation can determine desiccation resistance and whether we can identify the important CHC components that underlie variation in desiccation resistance across different species. Several large scale studies produced large datasets in determining the evolution of desiccation resistance across closely related *Drosophila* species (Kellermann *et al*., 2018; Kellermann *et al*., 2012; Matzkin et al., 2009) and ant species (Bujan et al., 2016; Hood and Tschinkel, 1990). Other studies produced large datasets in CHC variation across species (Khallaf et al., 2021; Menzel *et al*., 2017; Nunes et al., 2017; Van Oystaeyen et al., 2014) and focused on the communication aspects of CHCs. However, variations in CHCs and differences in desiccation resistance have not been experimentally connected in a phylogenetic framework. Analyzing CHC profiles and desiccation resistance across closely related species measured at the same condition can determine important CHC components that could predict various levels of desiccation resistance. Understanding the mechanisms underlying desiccation resistance and examining the evolution of these traits is key to predicting the species’ responses in facing drier environments that could result from future climate change (Chown et al., 2010).

In this study, we used a cohort-based experimental design of 50 *Drosophila* and related species, and experimentally determined their CHC compositions and desiccation resistance under similar conditions. Using a random forest machine learning algorithm, we built decision trees connecting CHC components and desiccation resistance in the 50 species and tested their correlation. First, we determined that CHC variations can largely explain differences in desiccation resistance across these species. Second, we identified a subset of CHCs, the methyl-branched CHCs (mbCHCs), as the most important CHC components that can determine desiccation resistance across these species. Third, we determined that the evolution of mbCHCs is also significantly correlated with evolution of higher desiccation resistance in *Drosophila* species. Given that mbCHCs are almost ubiquitous in insect species, we suggest that evolution of mbCHCs can be a general mechanism for insect species to evolve higher levels of desiccation resistance and adapt to more arid environments.

## Results

### A cohort-based experimental design for determining correlations between CHCs and desiccation resistance

To investigate how CHC variation across an evolutionary gradient affects desiccation resistance, for our experiments, we selected 46 *Drosophila* species representing both the *Sophophora* subgenus and the *Drosophila* subgenus, as well as three *Scaptodrosophila* species and one *Chymomyza* species, (**Figure S1**). As the assays for CHCs and desiccation resistance cannot be performed on the same individuals, we used a cohort-based experimental design (**Figure 1**) where subsets of each cohort (5-6 per species) were used for GC-MS determination of CHC profiles, desiccation resistance assays, and measuring body weight of the F1 progeny for each sex. Therefore, all measurements were performed at the cohort level.

**Figure 1.**
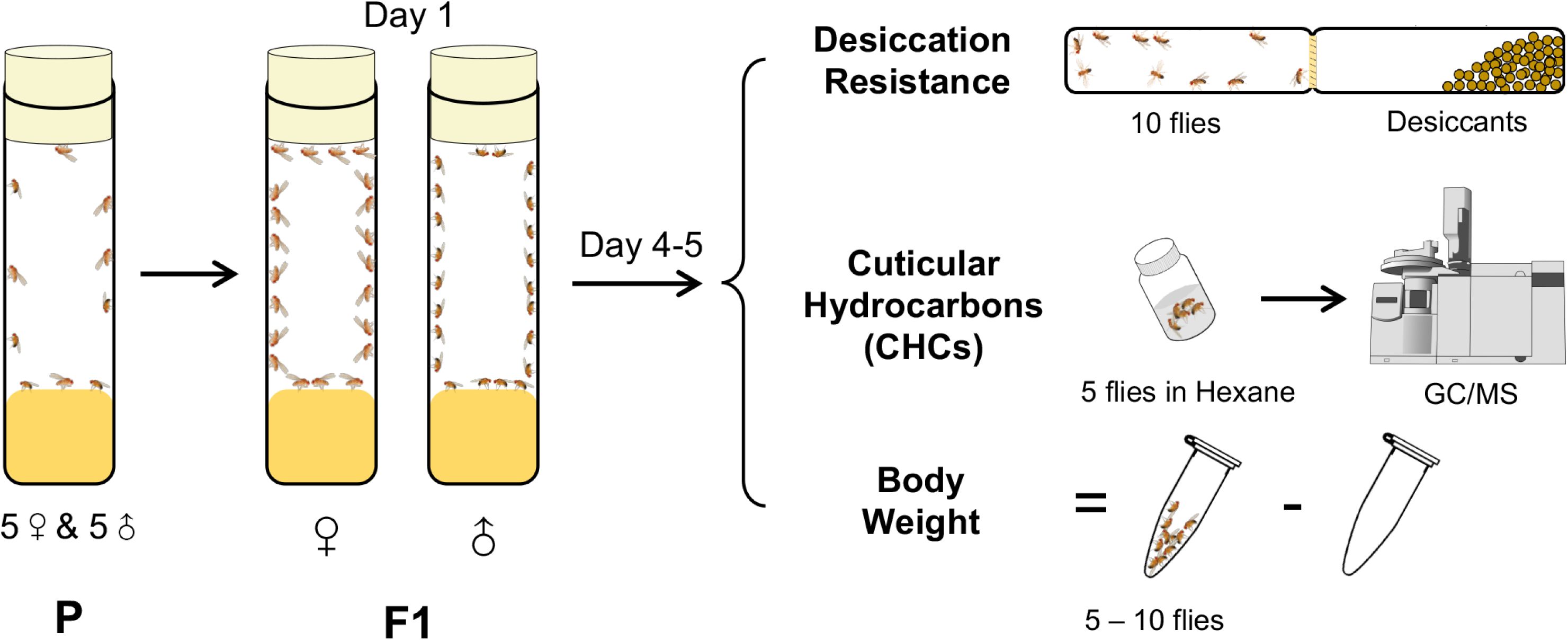
A cohort-based experimental design. A cohort-based experimental design can facilitate determining the correlation between desiccation resistance, cuticular hydrocarbons (CHCs), and body weight. Five to six cohorts of each species was established.

Desiccation resistance varied across these 50 species with desert-dwelling species from the repleta group showing the highest desiccation resistance (e.g., *D. mojavensis* males: 58.2 ± 4.7 hours) and species from melanogaster group showing the lowest desiccation resistance (e.g., *D. mauritiana* males: 2.7 ± 0.2 hours) (**Figure 2, Figure S2**), consistent with the findings of previous research (Kellermann *et al*., 2012; Matzkin *et al*., 2009). GC-MS analyses of the CHCs in these 50 species detected five types of CHCs with different carbon-chain lengths and quantities, including linear alkanes (*n*-alkanes), methyl-branched alkanes (mbCHCs; the methyl branch is on the second carbon), monoenes (with one double bond), dienes (with two double bonds), and trienes (with three double bonds) (**Figure S3**). *n*-alkanes, which have the highest melting temperatures, were only present in species from the melanogaster group, some of which have the lowest desiccation resistance among the species tested in this study. This suggests that *n*-alkanes may not make a general contribution to desiccation resistance across *Drosophila* species. Trienes, which have the lowest melting temperatures, were only present in eight species with low to moderate quantities. The other three types of CHCs, the mbCHCs, monoenes and dienes, were observed in most species tested in our study.

**Figure 2.**
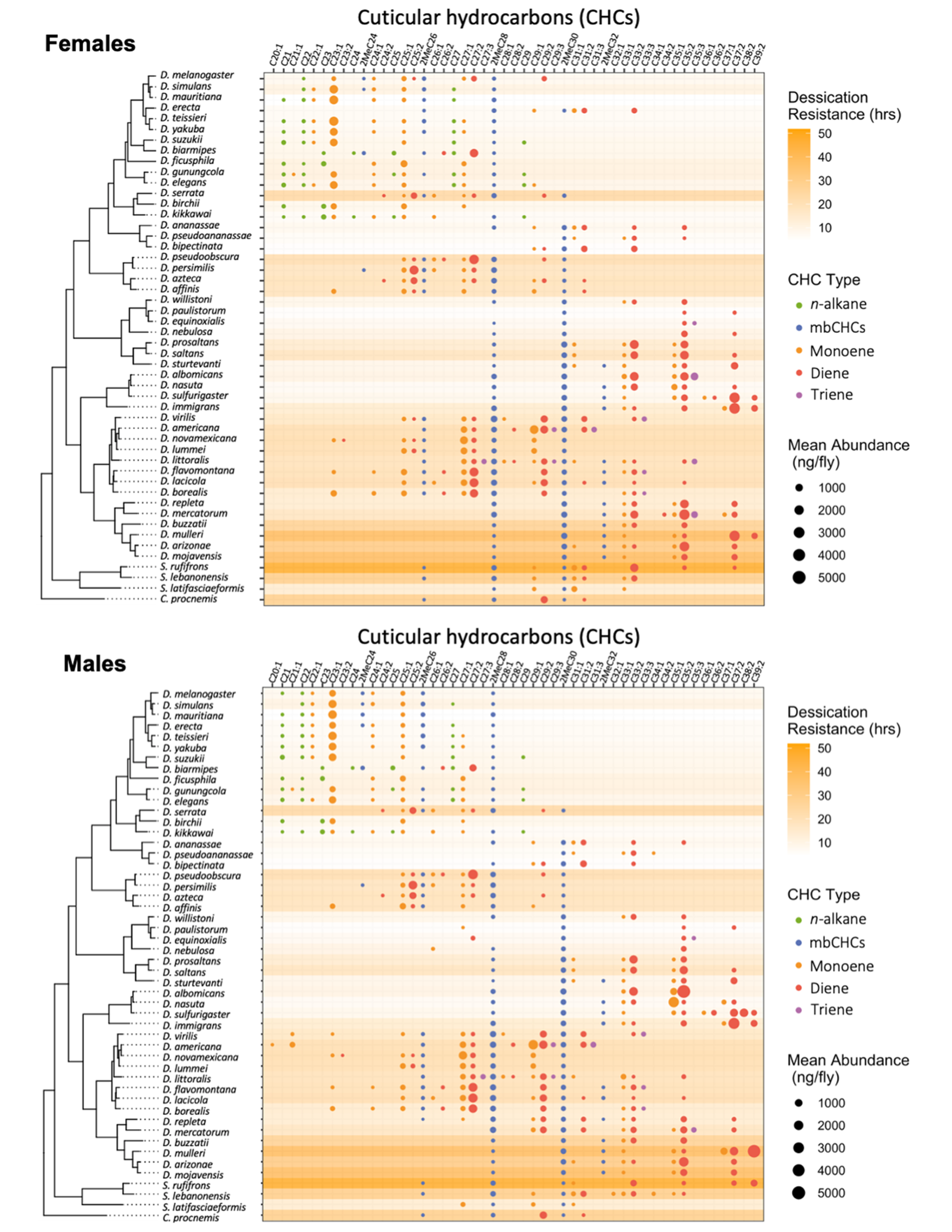
Desiccation resistance and CHC composition in 46 *Drosophila* species and 4 outgroup species. Desiccation resistance and CHC composition of each species were plotted together. Males and females are shown separately. The shading intensity of each species represents the number of hours of desiccation resistance, while the size and color of each circle represent the type of CHC and its quantity. *n*-alkanes are only present in some species from the melanogaster group.

### Higher relative quantities of CHCs are not primarily responsible for higher desiccation resistance

We first sought to determine if higher desiccation resistance in flies could be due to having higher amounts of CHCs. One caveat in the analysis between absolute CHC quantity and desiccation resistance is a possible correlation between CHC quantities and body size: species with larger body size may possess higher quantities of CHCs. This can lead to potential biases when directly determining correlations between CHC quantities and desiccation resistance. Species with higher body weight can also have higher absolute water content and a lower surface area to volume ratio due to their larger size, which may also lead to higher desiccation resistance (Wang *et al*., 2021).

Our initial analyses showed a significant positive correlation (Females: r = 0.4, *P* < 0.001; Males: r = 0.5, *P* < 0.001) between body weight and desiccation (**Figure 3A**). We also found a significant positive correlation between body weight and total amount of CHCs (Female: r = 0.7, *P* < 0.001; Male: r = 0.7, *P* < 0.001) (**Figure 3B**), suggesting that variation in the body weight (or size) of different species may be a confound in determining the relationship between CHC quantity and desiccation resistance. To take this confound into consideration, we normalized the total CHC quantity by dividing by the body weight for each species. This shows that CHCs account for 0.02 to 0.5% of the total body weight of different species (**Figure S4**). Analyses between the normalized CHC quantity and desiccation resistance showed no correlation between these two variables in males (Male: *P* = 0.1) and positive correlation in females (Female: r = 0.1, *P* = 0.03) (**Figure 3C**). The low value of r (0.1) indicates a weak correlation between CHC quantities and desiccation resistance in females. This suggests that having higher amounts of CHCs only has a limited contribution to higher desiccation resistance.

**Figure 3.**
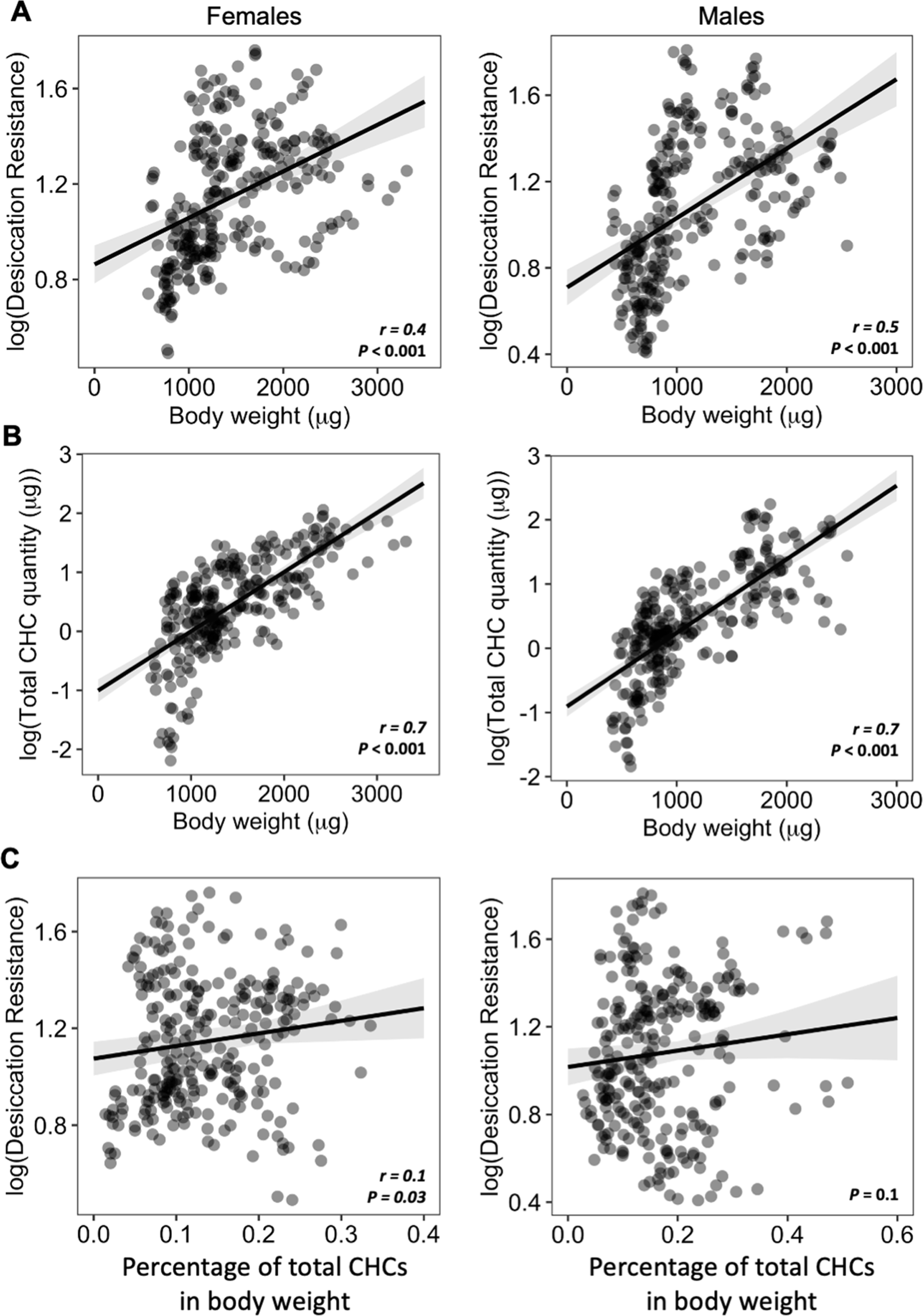
Higher amounts of CHCs do not contribute to higher desiccation resistance. **(A)** Body weight is positively correlated with desiccation resistance (Females: r = 0.4, *P* < 0.001, Males: r = 0.5, *P* < 0.001). **(B)** Total amount of CHCs is correlated with having higher body weight (Females: r = 0.7, *P* < 0.001, Males: r = 0.7, *P* < 0.001). **(C)** A weak positive correlation between desiccation resistance and CHCs as a percentage of body weight in females, while no correlation in males (Females: r = 0.1, *P* = 0.03, Males: *P* = 0.1). All correlation analyses were conducted using Pearson’s method.

### mbCHCs are important determinants of desiccation resistance

As total CHC quantity is not a main contributor to desiccation resistance in our study, we hypothesized that the composition of CHC profiles may be important for desiccation resistance. To test this hypothesis, we applied a permutational multivariate analysis of variance (PERMANOVA) to test whether the composition of CHCs differed across desiccation resistance (Anderson et al., 2006). The PERMANOVA analysis use beta diversity to represent differences in CHC compositions in these species and showed that the beta diversity of CHC compositions differs significantly across the increasing desiccation resistance (r^2^ = 0.1, *P* < 0.001; **Figure S5**), suggesting that CHC composition is important for desiccation resistance. We further sought to identify individual components of CHC profiles that are important determinants of desiccation resistance. To test if variation in CHC components can be used to determine desiccation resistance, we applied a random forest regression method that uses decision trees to connect the variables of interest (Liaw and Wiener, 2002; Svetnik et al., 2003), e.g., CHC composition and desiccation resistance, and identified the importance of individual CHC components to predict desiccation resistance. In this analysis, we treated the CHC profiles of each species and sex as an individual dataset, giving us 100 datasets (50 species x 2 sexes) with 5 to 6 individual CHC profiles each. We then correlated the decision trees that generated from the CHC composition with desiccation resistance. The prediction process can identify key CHC components that are important contributors to desiccation resistance.

Random forest modeling of CHC composition was able to explain 85.5% of the variation in desiccation resistance (Out of Bag Estimate: Root Mean Square Error ‘RMSE’ = 4.5) (**Figure 4A**). Models built with a 70:30 training/test split (n = 382, n = 164) performed similarly to models built with out of bag estimate (RMSE = 5.4, **Figure S6**). Four mbCHCs were identified as having the highest contribution to predicting desiccation resistance in the regression model (listed in decreasing importance: 2MeC30, 2MeC28, 2MeC32, and 2MeC26), while many other CHCs did not substantially contribute to predicting desiccation resistance (**Figure 4B**). These results suggest that CHCs, and mbCHCs in particular, are important predictors of desiccation resistance in *Drosophila* species.

**Figure 4.**
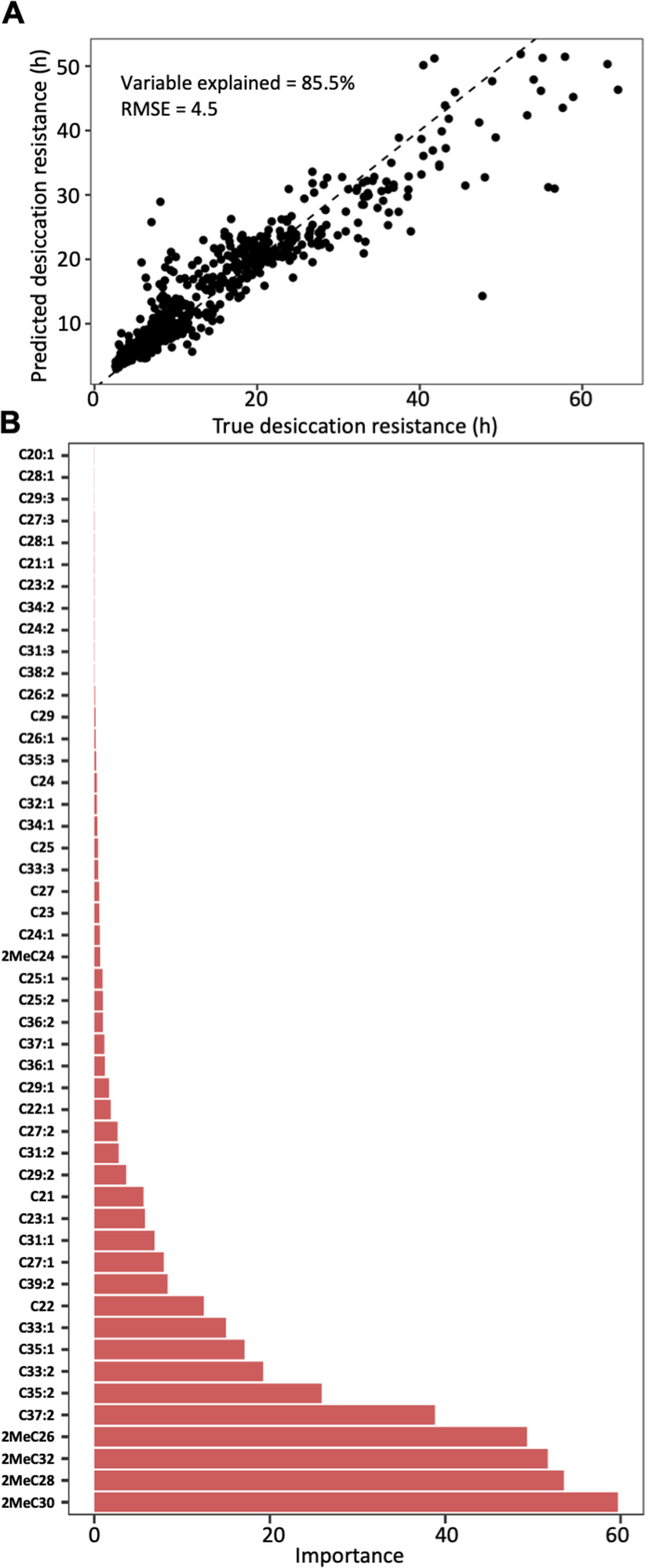
CHC composition can be used to predict desiccation resistance. **(A)** Random Forest regression modeling of CHC abundance was able to explain 85.5% of the variation in time to desiccation with a Root Mean Square Error (RMSE) of 4.5. **(B)** The abundance of four mbCHCs, 2MeC30, 2MeC28, 2MeC32, and 2MeC26 has the highest importance to the desiccation resistance in the random forest regression model, while most of CHCs have less contribution to the accuracy of the model for desiccation resistance.

### Longer mbCHCs confer higher desiccation resistance

We further sought to determine how mbCHCs contribute to desiccation resistance. As each species produces different combinations of these mbCHCs that are correlated with each other (**Figure S7**), we did not perform correlation analyses on these mbCHCs with a single regression model due to multicollinearity issues (Belsley et al., 2005). Instead, we performed correlation analyses between each of the most important CHCs identified from the random forest modeling, 2MeC26, 2MeC28, 2MeC30, and 2MeC32, and desiccation resistance. We found that the quantity of 2MeC26 negatively correlates with desiccation resistance in males (r = -0.4, *P* < 0.001) but no significant correlation was identified in females (*P* = 0.1) (**Figure 5A**). Significant positive correlations were observed between quantities of the longer mbCHCs and desiccation resistance for females (2MeC28: r = 0.4, *P* < 0.001; 2MeC30: r = 0.2, *P* = 0.009; 2MeC32: r = 0.3, *P* < 0.001) and males (2MeC28: r = 0.4, *P* < 0.001; 2MeC32: r = 0.4, *P* < 0.001) (**Figure 5B-D**).

**Figure 5.**
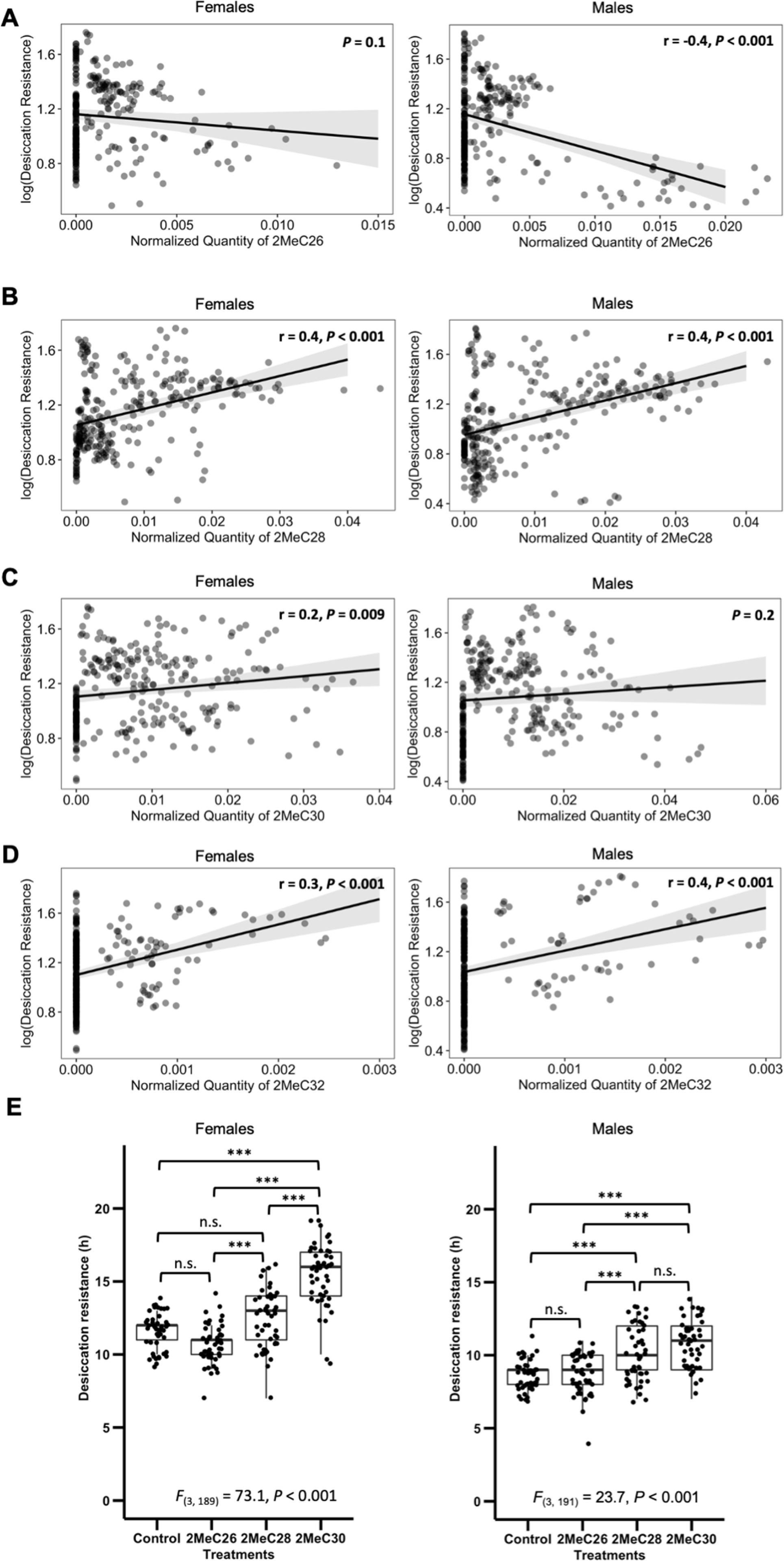
Longer mbCHCs are associated with higher desiccation resistance. **(A)** Quantity of 2MeC26 is negatively correlated with desiccation resistance in males, but no correlation in females (Females: *P* = 0.1, Males: r = -0.4, *P* < 0.001). **(B)** Quantity of 2MeC28 is positively correlated with desiccation resistance (Females: r = 0.4, *P* < 0.001, Males: r = 0.4, *P* < 0.001). **(C)** Quantity of 2MeC30 is positively correlated with desiccation resistance in females, but no correlation in males (Females: r = 0.2, *P* = 0.009, Males: *P* = 0.2). **(D)** Quantity of 2MeC32 is positively correlated with desiccation resistance (Females: r = 0.3, *P* < 0.001, Males: r = 0.4, *P* < 0.001). Correlations between the quantities of each mbCHC and desiccation resistance were determined using Pearson’s method. **(E)** *D. melanogaster* coated with longer mbCHCs have higher desiccation resistance. Perfuming of synthetic mbCHCs on *D. melanogaster* showed that 2MeC26 does not influence desiccation resistance. 2MeC30 increases desiccation resistance in female *D. melanogaster* while both 2MeC28 and 2MeC30 in increases desiccation resistance in *male D. melanogaster*. One-Way ANOVA showed significant differences between *D. melanogaster* flies coated with different mbCHCs (Female: *F*_(3,189)_ = 73.1, *P* < 0.001, Male: *F*_(3,191)_ = 23.7, *P* < 0.001). *Post hoc* comparison was conducted using Tukey’s method.

However, no significant correlation was observed between quantity of 2MeC30 and desiccation resistance in males (**Figure 5C**). These results suggested the production of longer mbCHCs could play a role in higher desiccation resistance.

To investigate the above results suggesting that longer mbCHCs could underlie higher desiccation resistance in *Drosophila* species, we performed desiccation assays on *D. melanogaster* coated with synthetic mbCHCs of different lengths (2MeC26, 2MeC28, and 2MeC30) at 25°C. While previous studies showed that a mixture of 2MeC26, 2MeC28, and 2MeC30 can rescue desiccation resistance in D. melanogaster flies without CHCs, experiments were not performed on individual mbCHCs for desiccation resistance (Krupp et al., 2020). Our desiccation assays showed that the coating of 2MeC26 did not increase desiccation resistance in both sexes (*post hoc* comparison with Tukey’s method followed by one-way ANOVA. Female: *P* = 0.07; Male: *P* = 1.0), while 2MeC30 significantly increased desiccation resistance in female *D. melanogaster* (*t* = 11.2, *P* <0.001), and both 2MeC28 (*t* = 4.6, *P* <0.001) and 2MeC30 (*t* = 6.9, *P* <0.001) significantly increased desiccation resistance in male *D. melanogaster* (**Figure 5E**). This suggests that production of longer mbCHCs can lead to higher desiccation resistance.

### Evolution of mbCHCs underlies variation in desiccation resistance in *Drosophila* species

We have shown that mbCHCs are important determinants of desiccation resistance in *Drosophila* species and experimentally coating insects with longer mbCHCs confers higher desiccation resistance. To further determine whether the evolution of mbCHCs underlies the variation in desiccation resistance in *Drosophila* species, we first examined the evolutionary trajectory of mbCHCs and then tested the correlation between mbCHCs and desiccation resistance with phylogenetic correction. We mapped the evolution of mbCHC composition in the *Drosophila* genus using ancestral trait reconstruction and tested the phylogenetic signal using Pagel’s *λ* (Freckleton et al., 2015; Goolsby et al., 2017). Ranging from 0 to 1, Pagel’s *λ* measures the extent to which the variance of the trait can be explained by the phylogeny and therefore determines the degree of association between the trait evolution and the phylogeny (Lynch, 1991). If *λ* is closer to 1, the evolution of the trait has a higher association with the phylogeny. Phylogenetic signals in mbCHCs were detected for both females and males with *λ* = 0.75 and 0.83, respectively (**Table S1**). This suggests a moderate to strong association between mbCHC evolution and phylogeny. The derived ancestral state for mbCHCs in the *Drosophila* genus suggest that 2MeC28 and 2MeC30 are the major mbCHCs. (**Figure S8**). The evolutionary trajectory of mbCHC composition showed repeated evolution in the lengths of mbCHCs, as well as their quantities. Independent evolution of longer mbCHCs was observed in four clades of species, including the willistoni, nasuta, and repleta groups, and the ananassae subgroup (including *D. ananassae*, *D. pseudoananassae*, and *D. bipectinata*), while shorter mbCHCs were mainly observed in several species in the melanogaster subgroup (e.g., *D. melanogaster*, *D. simulans*, and *D. teissieri*) (**Figure 2 & S8**).

Desiccation resistance also has a strong association with the phylogeny of *Drosophila* species (**Table S2)** (Kellermann *et al*., 2018; Kellermann *et al*., 2012). We sought to determine if the variation of desiccation resistance could be explained by the evolution of mbCHCs in these species. We used a Phylogenetic Generalized Linear Square (PGLS) model to test for the correlation between mbCHCs and desiccation resistance when controlling the effects from the phylogenetic relationship of the species (Grafen, 1989; Mundry, 2014). Since mbCHCs are produced in a linear pathway with each species producing several different mbCHCs, we selected the mbCHC with the longest carbon-chain length in the species as the proxy to represent the lengths of mbCHCs in each species. We incorporated the quantity, length, and their interaction in the PGLS model for the longest mbCHC. The interaction term between the length and quantity of the longest mbCHC in the model can determine how the two variables combinatorically affect desiccation resistance. PGLS modeling showed, after correcting for phylogenetic effects, the higher quantity and longer length of the longest mbCHC both affect desiccation resistance (interaction term, Female: *t* = 3.5, *P* < 0.001; Male: *t* = 2.2, *P* = 0.03) (**Table S3; Figure 6**). This suggests that the synthesis of mbCHCs with longer carbon-chain lengths could be a common mechanism underlying the evolution of higher desiccation resistance.

**Figure 6.**
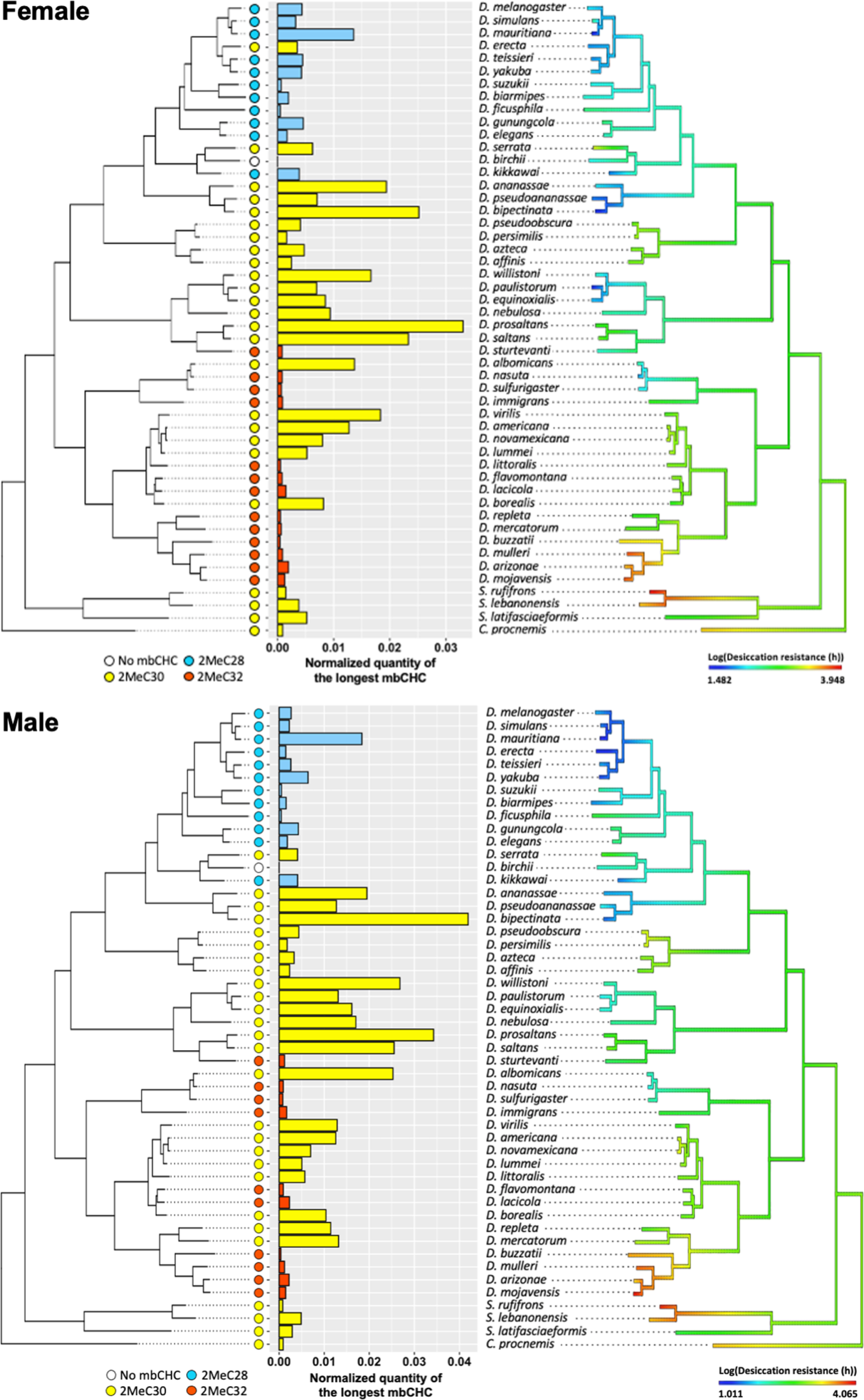
Evolution of longer mbCHCs are significantly correlated with evolution of higher desiccation resistance. Patterns in the normalized quantities of the longest mbCHCs and desiccation resistance for females (top) and males (bottom) were listed across the phylogeny of the 50 *Drosophila* and related species. PGLS analysis between the longest mbCHCs and desiccation resistance showed both the higher quantity and longer length of the longest mbCHC affect desiccation resistance (interaction term, Female: *t* = 3.5, *P* < 0.001; Male: *t* = 2.2, *P* = 0.03).

## Discussion

Reducing evaporative water loss through the cuticle using a layer of CHCs is one of the most important evolutionary innovations in insects that allows many species to survive and thrive in diverse and arid habitats. We showed that CHC composition can account for 85.5% of the variation in desiccation resistance in the 50 *Drosophila* and related species in this study. This suggests that CHC composition may be highly predictive of desiccation resistance. Algorithmic ranking in importance of CHC components using a random forest machine learning model showed that mbCHCs have the highest contribution to determining desiccation resistance. Importantly, higher amounts of longer mbCHCs are important in desiccation resistance. This is consistent with previous studies showing that mbCHC has higher melting temperature than the other commonly present types of CHCs (monoene and diene) and longer CHC leads to higher melting temperature (Gibbs and Pomonis, 1995). This is also experimentally supported by the synthetic CHC coating experiments where coating of longer mbCHCs confers higher desiccation resistance.

Previous studies showed support for mbCHCs in desiccation resistance in *Drosophila*. RNAi knockdown of a methyl-branched specific fatty acid synthase (*mFAS*) in *D. serrata* eliminates almost all mbCHCs and leads to significant decrease in desiccation resistance which could be partially rescued with synthetic mbCHCs (Chung *et al*., 2014). Its sibling species, the rainforest desiccation sensitive *D. birchii*, have only trace amounts of mbCHCs (Howard et al., 2003) and is unable to evolve desiccation resistance despite strong laboratory selection over many generations (Hoffmann et al., 2003). The presence of mbCHCs in most *Drosophila* and related species leads to another question: Is this a general rule for insects using mbCHCs to develop desiccation resistance? A high proportion of very-long-chain mbCHCs with a single methyl branch have been reported in many insect species from different taxa that dwell in desert environments, such as the desert tenebrionid beetle *Eleodes armata* (Hadley, 1977), the desert ants *Cataglyphis niger* and *Pogonomyrmex barbatus* (Johnson and Gibbs, 2004; Soroker and Hefetz, 2000), and the desert locust *Schistocerca gregaria* (Heifetz et al., 1998). Although a small proportion of *n-* alkanes was also reported in some of these species, this suggests the use of mbCHCs to minimize water evaporation may be a common mechanism for insects in extremely dry environments.

Why are *n*-alkanes, which have the highest melting temperature and potentially better water proofing properties, not prevalent in species in our studies? A hypothesis that could explain this is the competition on the common precursors between the synthesis of linear unsaturated CHCs (monoenes and dienes) and *n*-alkanes. In insects, the synthesis of all CHCs start from the fatty acyl CoA synthesis pathway. The pathway is split where a cytosolic Fatty Acid Synthase (cFAS) will synthesize all the linear CHCs (alkanes, monoenes, dienes), while mFAS will synthesize mbCHCs (**Figure S9**) (Chung and Carroll, 2015; Holze et al., 2021). The pathway suggests that the synthesis of alkanes, monoenes, and dienes are in competition. As many monoenes and dienes function in *Drosophila* as contact pheromones in regulating different behaviors such as mating and aggression (Blomquist and Ginzel, 2021; Chertemps et al., 2007; Chertemps et al., 2006; Chung and Carroll, 2015; Krupp et al., 2008; Wang et al., 2011), the synthesis for these CHCs may compete with the synthesis of *n*-alkanes. In contrast, the synthesis of mbCHCs is in another part of the pathway. This suggests that there are potential constraints in the roles of *n*-alkanes in desiccation resistance in *Drosophila* due to competition with CHCs which function as signaling molecules, while the synthesis of mbCHCs is relatively unaffected by other products of the pathway (**Figure S10**) (Chung and Carroll, 2015). We speculate that in *Drosophila* and related species, the use of mbCHCs for modulating desiccation resistance may avoid the conflict between surviving desiccation and chemical signaling.

If there are no biosynthetic constraints to the synthesis of mbCHCs, would the evolution of longer mbCHCs be a general mechanism in the evolution of higher desiccation for species adapting to more arid climates? We suggest that this may be the case for many species, but for some species that use the shorter mbCHCs such as 2MeC26 and 2MeC28 as signaling molecules, such as *D. serrata* (Chenoweth and Blows, 2005; Chung *et al*., 2014) and the longhorned beetle, *Mallodon dasystomus* (Spikes et al., 2010), natural selection and sexual selection may have opposing effects on the lengths of these mbCHCs. Indeed, the evolution of longer CHCs can have a sexually antagonistic effect on desiccation and mating success (Rusuwa et al., 2022).

Ancestral trait reconstruction showed that the derived ancestral status for mbCHCs in the *Drosophila* genus are 2MeC28 and 2MeC30 as the major mbCHCs (**Figure S8**). As there are many extant species that do not have 2MeC30, this suggests that the ability to synthesize longer mbCHCs may be lost during evolution, especially for species that do not inhabit arid environments and have low levels of desiccation resistance such as *D. melanogaster* and *D. biarmipes*. On the contrary, for species that dwell in extremely arid environments such as *D. mojavensis* and *D. arizonae*, higher quantities of 2MeC30 and an even longer mbCHC, 2MeC32, are observed. The observation is consistent with the results from our PGLS modeling, showing significant correlation between having higher quantity of longer mbCHCs and higher desiccation resistance in the *Drosophila* genus. In summary, we suggest that the synthesis of mbCHCs with longer carbon-chain lengths could be a common mechanism underlying the evolution of higher desiccation resistance.

## Materials and Methods

### Drosophila species

In this study, 46 *Drosophila* species, as well as three *Scaptodrosophila* species and one *Chymomyza* species were either obtained from the National Drosophila Species Stock Center (NDSSC) or were gifts from various colleagues. Details are listed in **Table S4**. All species were reared on standard cornmeal medium (Flystuff 66-121 Nutri-Fly^®^ Bloomington Formulation). The phylogeny of all 50 species in this study was obtained from Finet et al. (2021).

### Experimental design

To investigate the contribution of cuticular hydrocarbons to desiccation resistance, a cohort-based design was used. For each species, five to six cohorts were established and three measurements were conducted for each sex of the F1 progeny, including desiccation resistance, cuticular hydrocarbons, and body weight (**Figure 1**). Each cohort was treated as a biological replicate. Each cohort in each species was established by pooling five females and five males on a standard cornmeal medium in the environmental chambers set at 25°C and a 12L:12D photoperiod. To maximize the food and spatial availability and minimize competition between the F1 progenies (Mueller, 1988), the parent flies in each cohort were transferred to fresh food after 5 days. F1 progeny flies were collected daily following emergence, separated by sex, and maintained on fresh cornmeal medium. All flies used for measurements were four- to five-days-old.

### Desiccation resistance assays

Desiccation resistance assays were performed in a randomized and blinded manner with the setup consistent with a previously published protocol (**Figure 1**) (Chung *et al*., 2014). Briefly, in each cohort, ten adults of the same sex were subjected to the assay setup containing 10 grams of silica gel (S7500-1KG, Sigma-Aldrich^®^, St. Louis, MO). After the assays were assembled, all the setups were randomly arranged with a number assigned. Mortality of the flies was recorded hourly after two hours. For each cohort, one vial was scored and the average time in hours until all flies died was recorded as desiccation resistance. All desiccation resistance assays were conducted in the same environmental chambers that rearing the flies at 25°C and 12L:12D photoperiod.

### Cuticular hydrocarbon analyses

Cuticular hydrocarbons (CHCs) were extracted and analyzed using GC-MS following previously published protocols (Lamb et al., 2020; Savage *et al*., 2021). Five flies of the same sex (four- to five-days-old) from each cohort were soaked for 10 min in 200 µl hexane containing hexacosane (C26; 25 ng/ul) as an internal standard. Extracts were directly analyzed by GC-MS (7890A, Agilent Technologies Inc., Santa Clara, CA) using a DB-17ht column (Agilent Technologies Inc., Santa Clara, CA). To identify the CHC composition, we first compared retention times and mass spectra to an authentic standard mixture (C7-C40) (Supelco^®^ 49452-U, Sigma-Aldrich, St. Louis, MO) with CHC samples. The types of CHCs, including methyl-branched alkanes, monoenes, dienes, and trienes, were then identified by a combination of their specific fragment ions on the side of functional groups (methyl branch or double bonds), retention times relative to linear hydrocarbon standards, and the m/z value of the molecular ion. The position of methyl branch in mbCHCs was determined using the protocol described in Carlson et al. (1998), while the position of double bonds were not determined in this study. Each CHC peak was quantified using its comparison with the peak area of the internal standard (C26) and represented as nanogram per fly (ng/fly). Because we observed a biased integration of peak areas on longer CHCs in running the standard mixture (**Figure S10**), we corrected the peak areas of CHCs based on the carbon-chain lengths using the integration from the standard mixture.

### Body weight measurement

Body weight was incorporated in the data collection and analysis. The body weight was determined as the difference between the Eppendorf tube containing five to ten flies of the same sex and the same empty Eppendorf tube.

### Coating of synthetic compounds

To coat each of synthetic mbCHCs on *D. melanogaster*, including 2MeC26, 2MeC28, and 2MeC30, we first added 200 µL hexane containing 300 ng/µL of each mbCHC in a 2-mL glass vial and then used a nitrogen evaporator (BT1603 G-Biosciences^®^) to evaporate the hexane with only mbCHCs precipitate at the bottom of the vial. For the control group, we only added 200 µL hexane without any mbCHC. After the hexane was evaporated, ten flies of the same sex were transferred into each vial, following shaking on a Vortex for 20 seconds on, 20 seconds off, and 20 seconds on. The flies were then directly subjected to desiccation assays. Synthetic mbCHCs were kindly provided by Dr. Jocelyn Millar (University of California, Riverside).

### Statistics

All analyses were conducted in R (Version 4.1). Correlation analyses were conducted with Pearson’s method using ‘*cor.test*’ function. The dependent variables were log-transformed to better conform with assumptions of normality. Variance in CHC beta diversity across desiccation resistance was determined using PERMANOVA using the ‘vegan’ package (Anderson *et al*., 2006). The random forest regression analysis was performed using the ‘ranger’ and ‘randomForest’ packages (Liaw and Wiener, 2002; Wright and Ziegler, 2015). The random forest regression models were built using both Out Of Bag estimate and test/training sets (70:30 split). To determine how useful each CHC variable is in the prediction of desiccation resistance in the random forest regression analysis, the importance of top predictor CHCs was quantified using permutation importance (Altmann et al., 2010). Desiccation resistance in flies coated with different mbCHCs were determined using one-way ANOVA at *alpha* = 0.05. *Post hoc* comparison was further conducted using Tukey’s method. The ancestral trait reconstruction and estimation of phylogenetic signal, Pagel’s *λ*, for mbCHC composition and desiccation resistance was determined using ‘Phytools’, ‘Picante’, and ‘Rphylopars’ packages (Goolsby *et al*., 2017; Kembel et al., 2010; Revell, 2012). The PGLS analysis was conducted using generalized least squares fit by maximum likelihood and the covariance structure between species was used under a Brownian motion process of evolution. The PGLS analyses were conducted using ‘*GLS’* function in ‘*ape’* package (Paradis and Schliep, 2019).

## ACKNOWLEDGEMENTS

We thank Dr. Jocelyn Millar for guidance and advice, as well as the synthetic compounds used in this study. We also thank Ye Ma, Eliana Giannetti, and Taylor Hori for technical assistance, Yushan Zhang for assistance with figure visualization, and the National Drosophila Species Stock Center for fly stocks of these different *Drosophila* species. This work is supported by a National Science Foundation grant (2054773) to H. Chung.

## AUTHOR CONTRIBUTIONS

Z.W. and H. Chung designed research; Z.W., J.P., H. Cong, and C.R. performed research; Z.W., J.R., and M.L. analyzed data; Z.W. and H. Chung wrote the paper with input from other authors.

## DATA ACCESSIBILITY

Data and code have been deposited in the Dryad Digital Repository at:

## DECLARATION OF INTERESTS

The authors declare no competing interests.

## Supplementary Data

### SUPPLEMENTARY FIGURE LEGENDS

**Figure S1.**
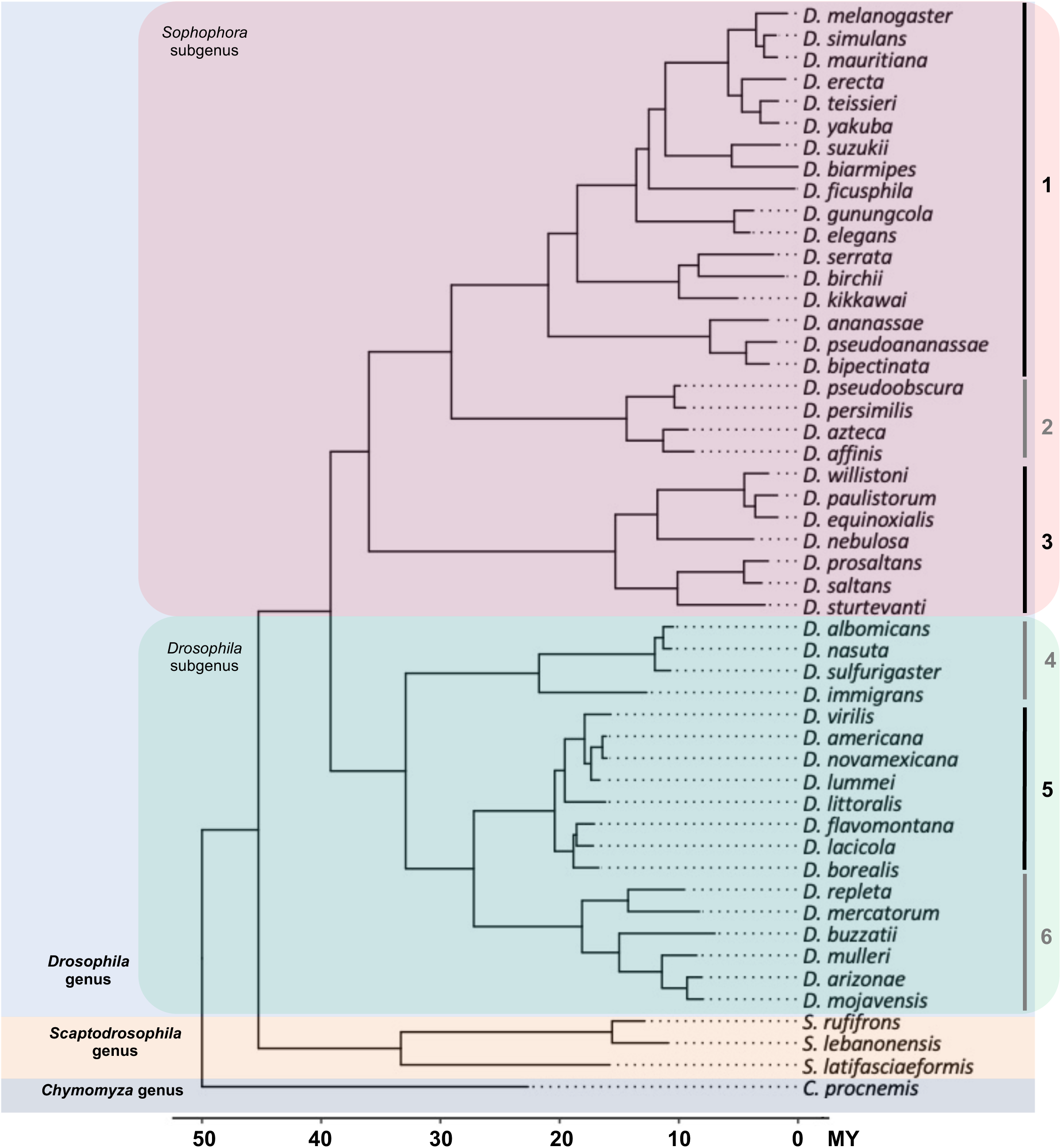
The phylogeny of 50 *Drosophila* and related species used in this study. 50 species representing 46 *Drosophila* species (18 from *Drosophila* subgenus, 28 from *Sophophora* subgenus) and four species (3 *Scaptodrosophila* species and 1 *Chymomyza* species) as outgroup were used in our study. The 46 *Drosophila* species represents some of the main groups in the *Drosophila* genus: (1) melanogaster group, (2) obscura group, (3) willistoni group, (4) nasuta group, (5) virilis group, and (6) repleta group.

**Figure S2.**
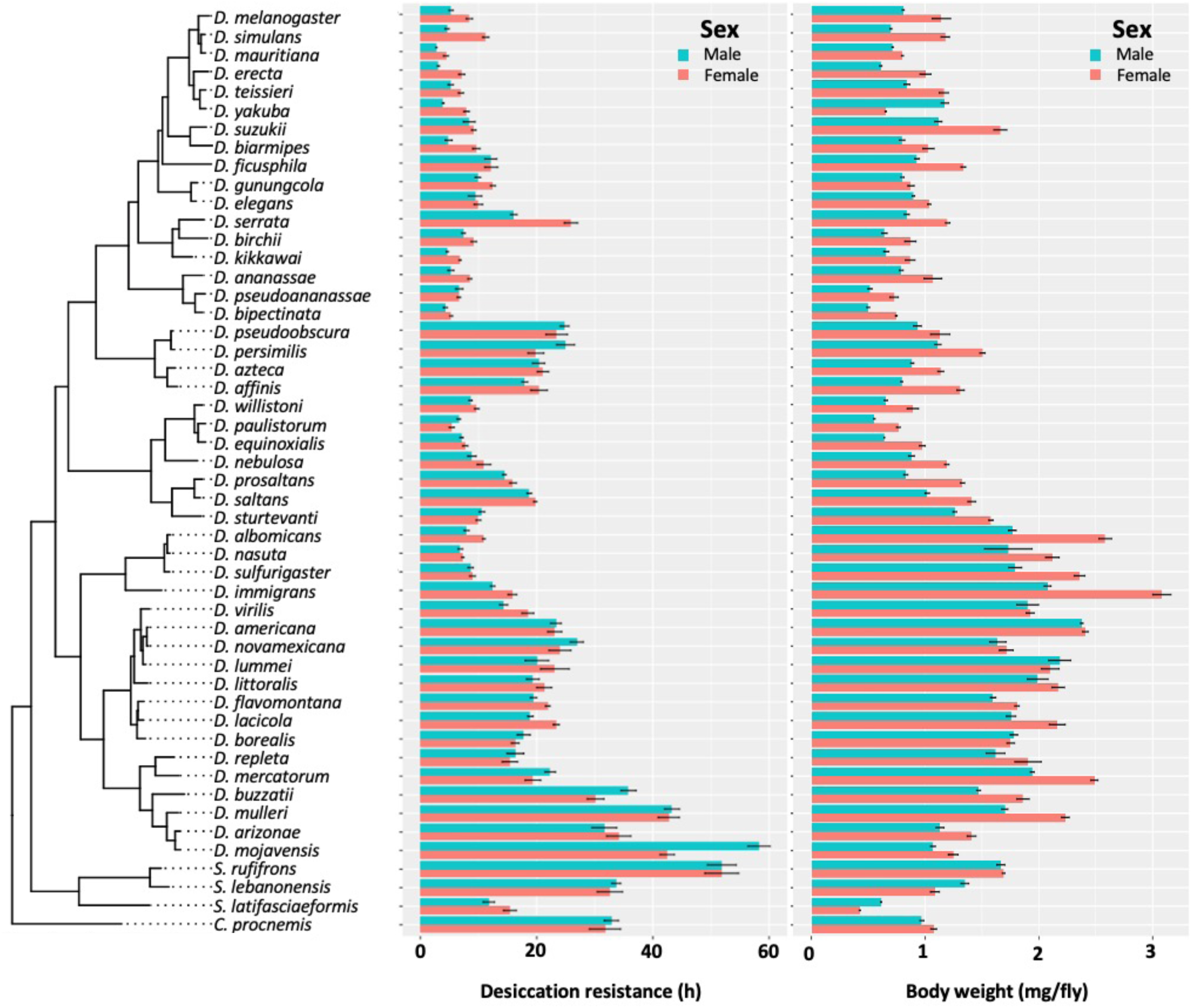
Desiccation resistance and body weight in 46 *Drosophila* species and 4 outgroup species. Desiccation resistance and body weight are shown in separate graphs.

**Figure S3.**
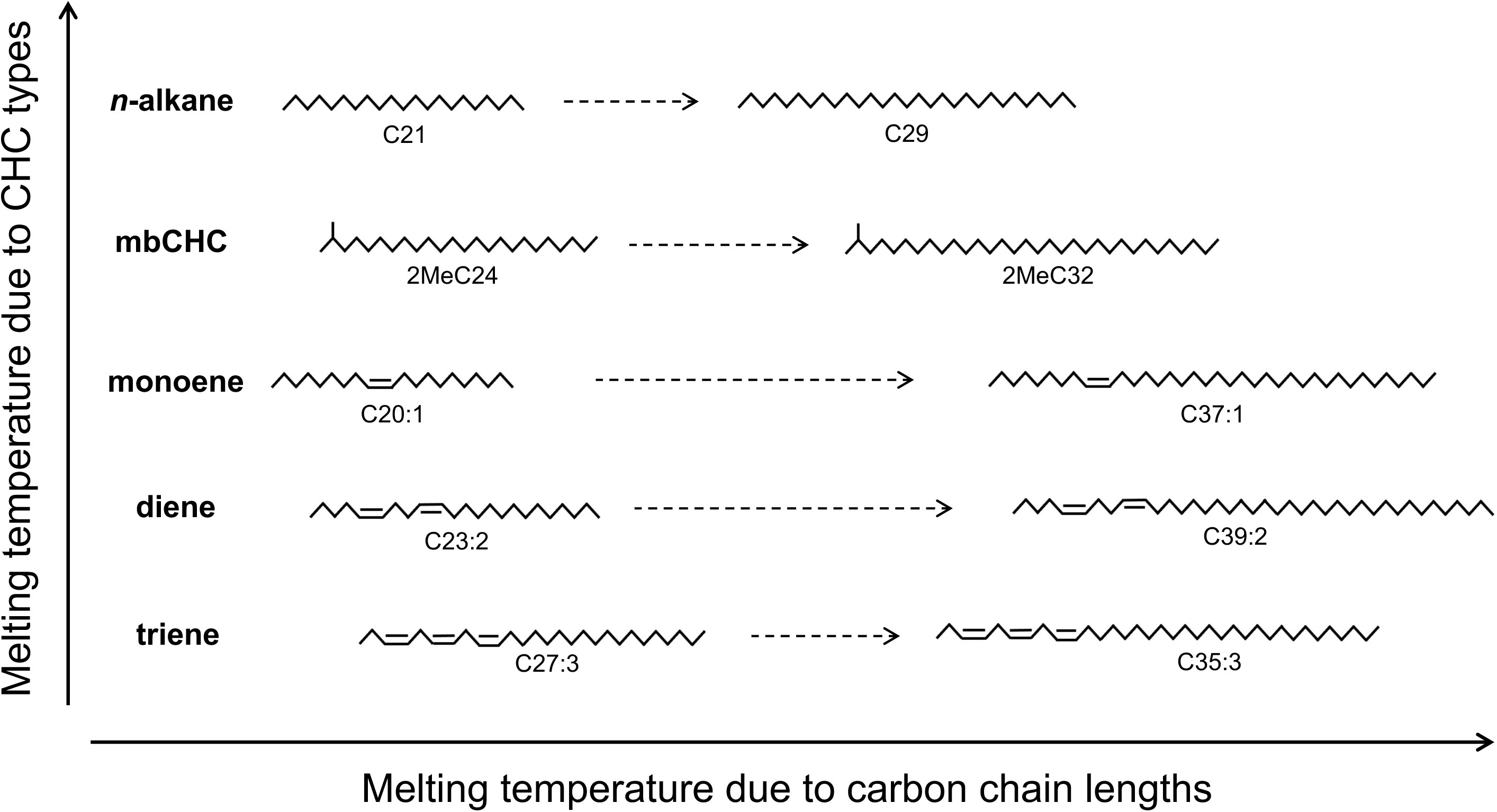
The melting temperature of CHCs is determined by methyl group, double bonds, and carbon-chain length. Five types of CHCs are detected in this study. Due to their chemical structures, *n*-alkanes have the highest melting temperatures followed by mbCHCs, monoenes, dienes, and trienes. Increasing length of the carbon-chain can also increase melting temperature. The positions of double bonds in this figure are for illustrative purposes and were not determined in this study.

**Figure S4.**
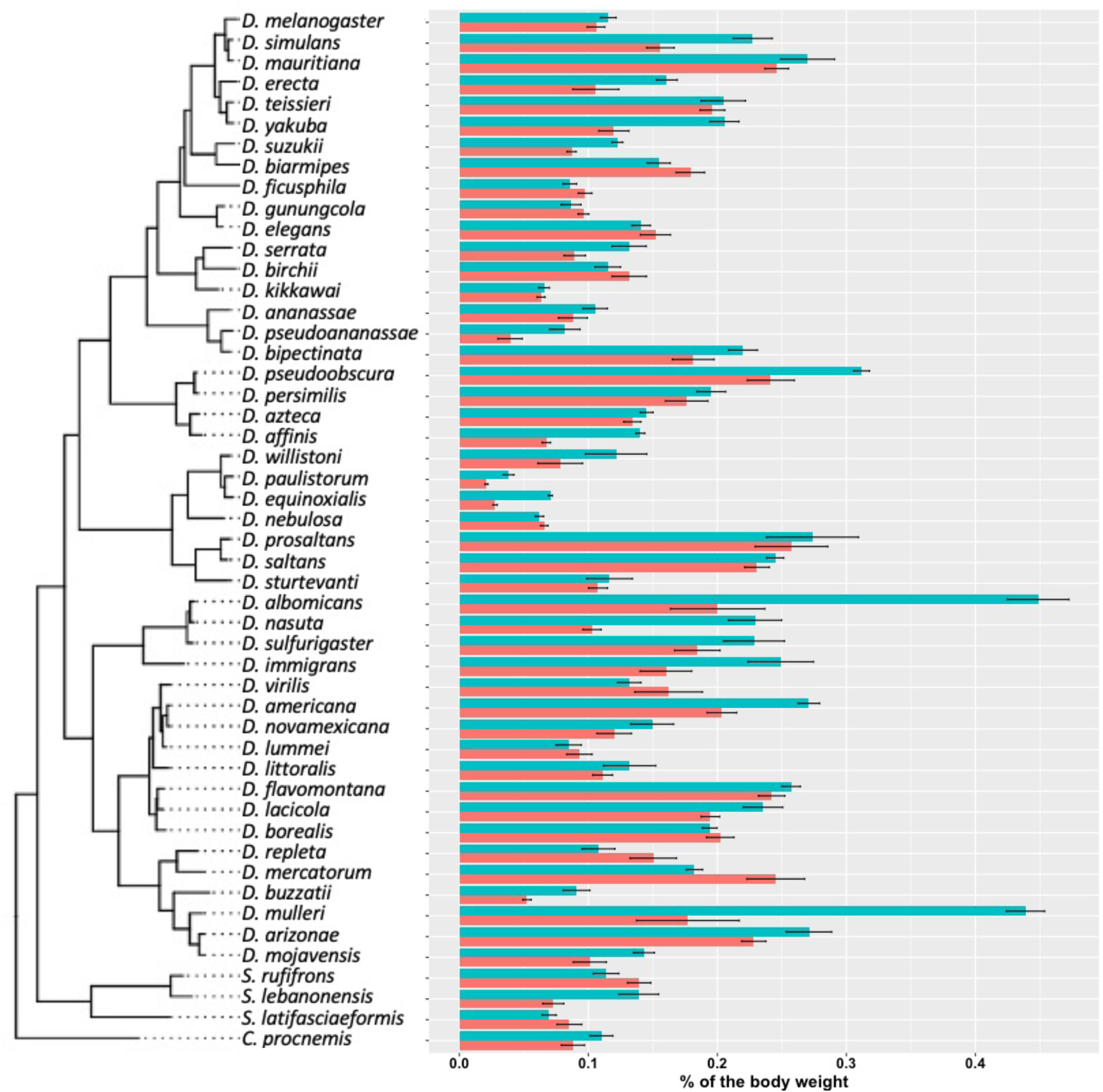
CHC quantity as a percentage of body weight. As *Drosophila* species largely vary in size, we normalized the total CHCs as a percentage of body weight. CHCs account from 0.02% to 0.5% of the total body weight of different species.

**Figure S5.**
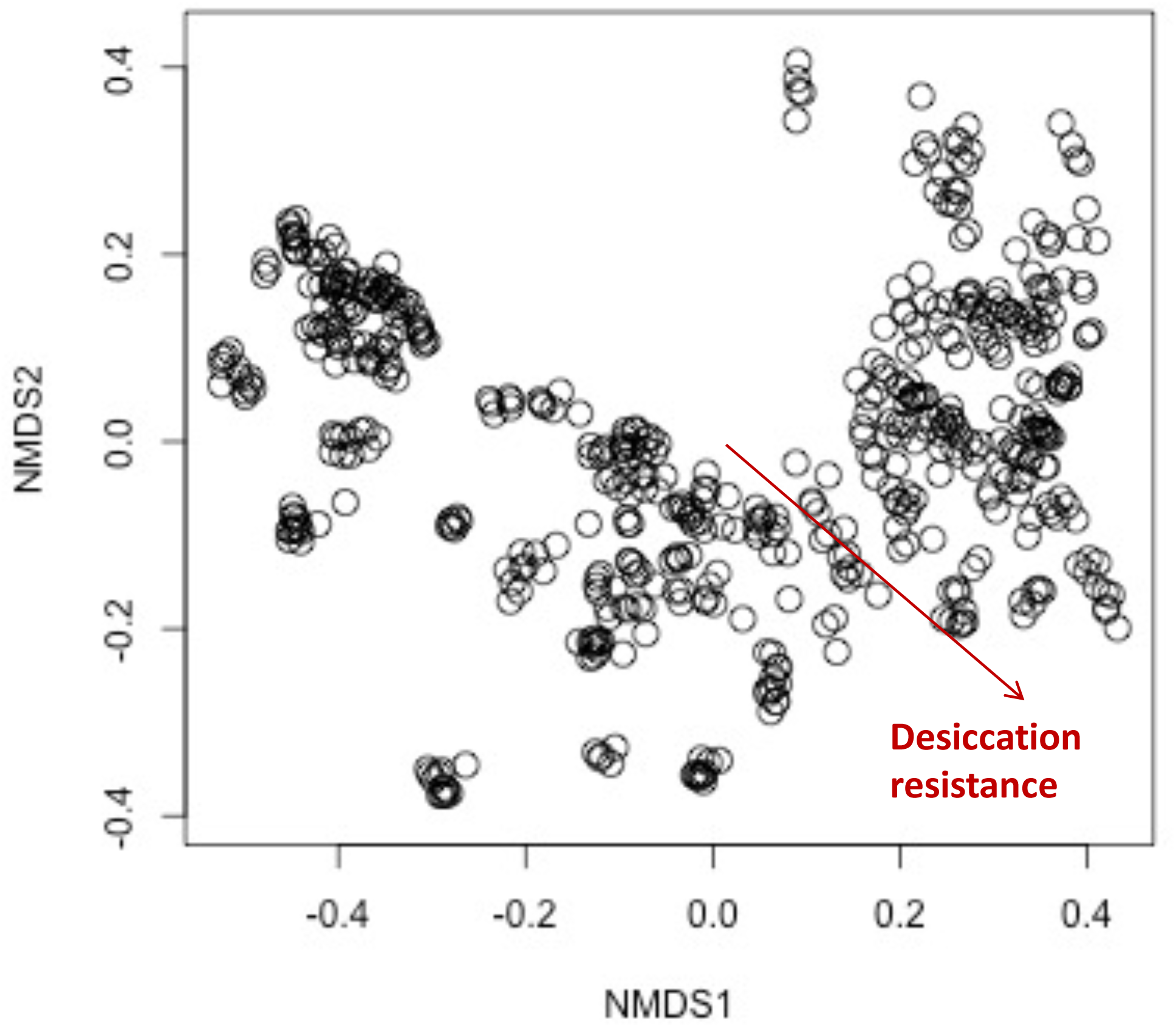
The composition of CHCs significantly differed across the increasing desiccation resistance. NMDS plot showing the beta diversity of CHCs differs significantly across the increasing desiccation resistance (PERMANOVA: r^2^ = 0.1, *P* < 0.001), suggesting CHC composition could affect desiccation resistance in the 50 *Drosophila* and selected species.

**Figure S6.**
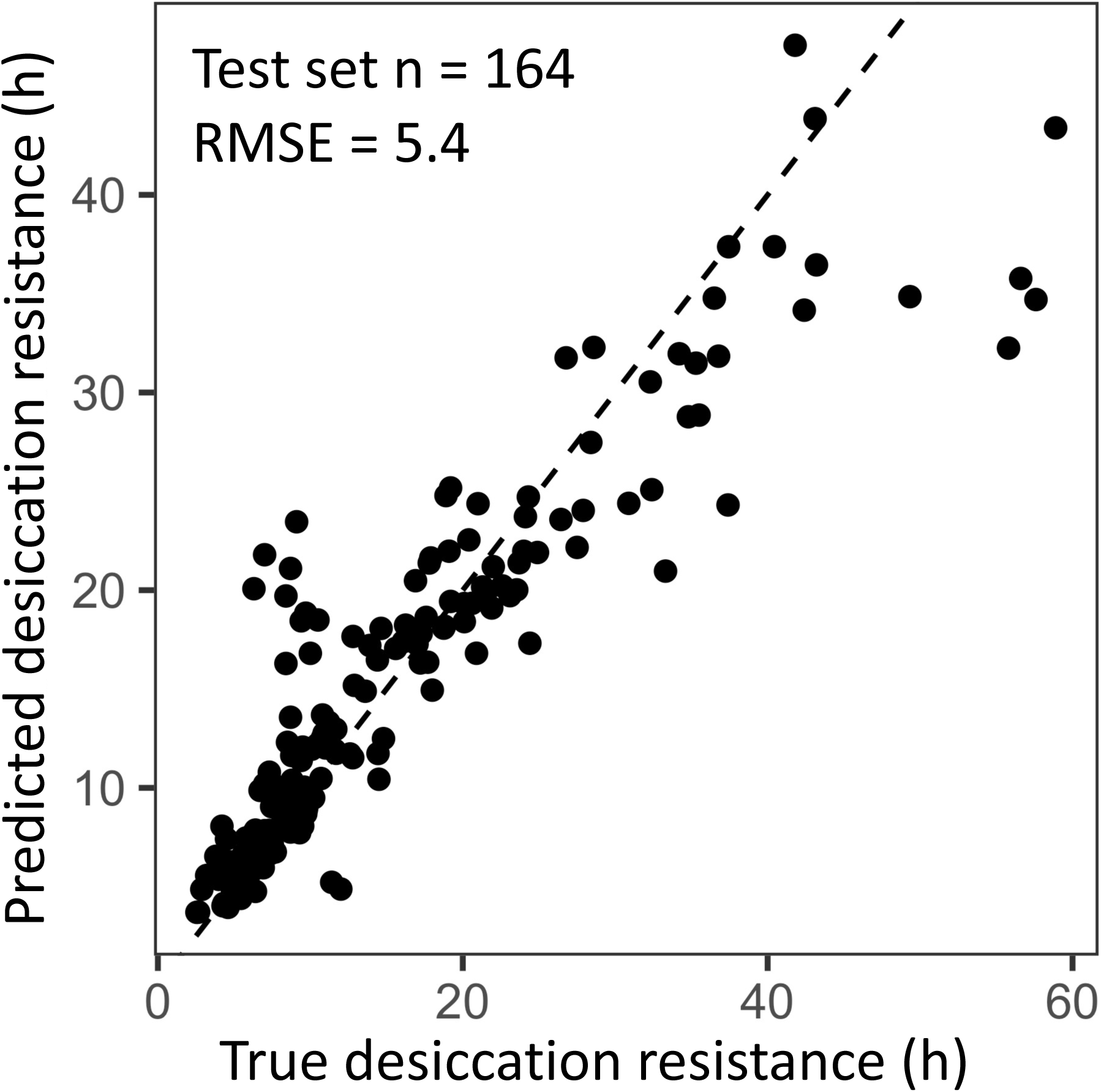
Cross-validation of the random forest regression model has a similar performance to the regression model using the full dataset. A 70:30 training/test split in the dataset (n = 382, n = 164) was used to evaluate the performance of the random forest regression model. The out of bag estimate of the root mean square error (RMSE) in the split dataset is 5.4, which is similar to the RMSE using the full dataset (RNMSE = 4.5).

**Figure S7.**
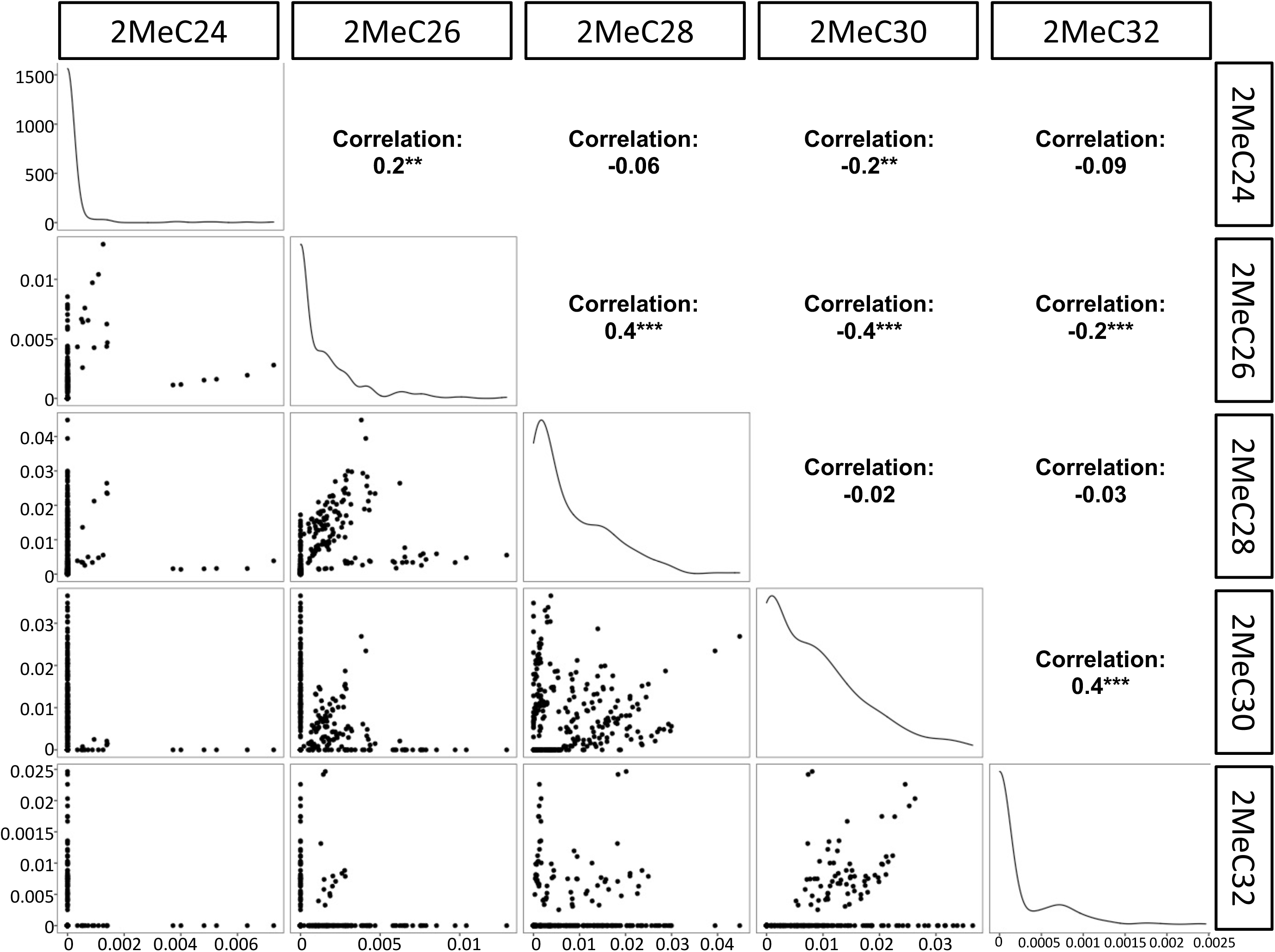
Correlation between different mbCHCs in 50 *Drosophila* and related species. Correlation between different mbCHCs was determined using Pearson’s method. Significant correlations between each pairs of mbCHCs were labeled with stars following with the correlation coefficients. **: *P* < 0.01; ***: *P* < 0.001.

**Figure S8.**
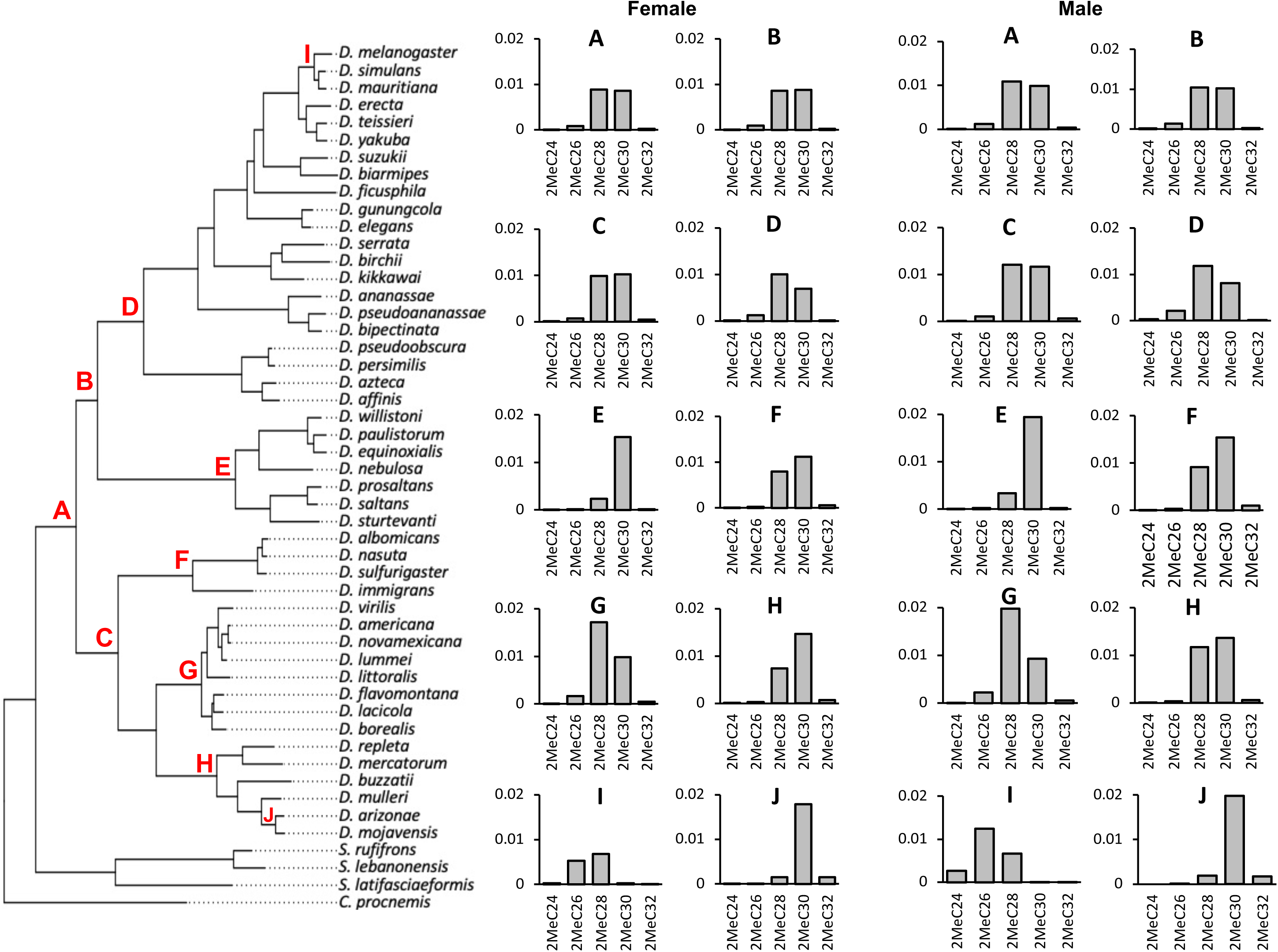
Ancestral state reconstruction of mbCHC for the *Drosophila* genus. Bar plots on the right were the derived ancestral state of female and male mbCHC profiles for the common ancestor of the *Drosophila* genus (A), the *Sophophora* subgenus (B), the *Drosophila* subgenus (C), the melanogaster group (D), the willistoni group (E), the nasuta group (F), the virilis group (G), the repleta group (H), the clade of *D. melanogaster*, *D. simulans*, and *D. mauritiana* (I), and the clade of *D. mojavensis* and *D. arizonae* (J). 2MeC28 and 2MeC30 were the major CHCs in the derived ancestral state (A).

**Figure S9.**
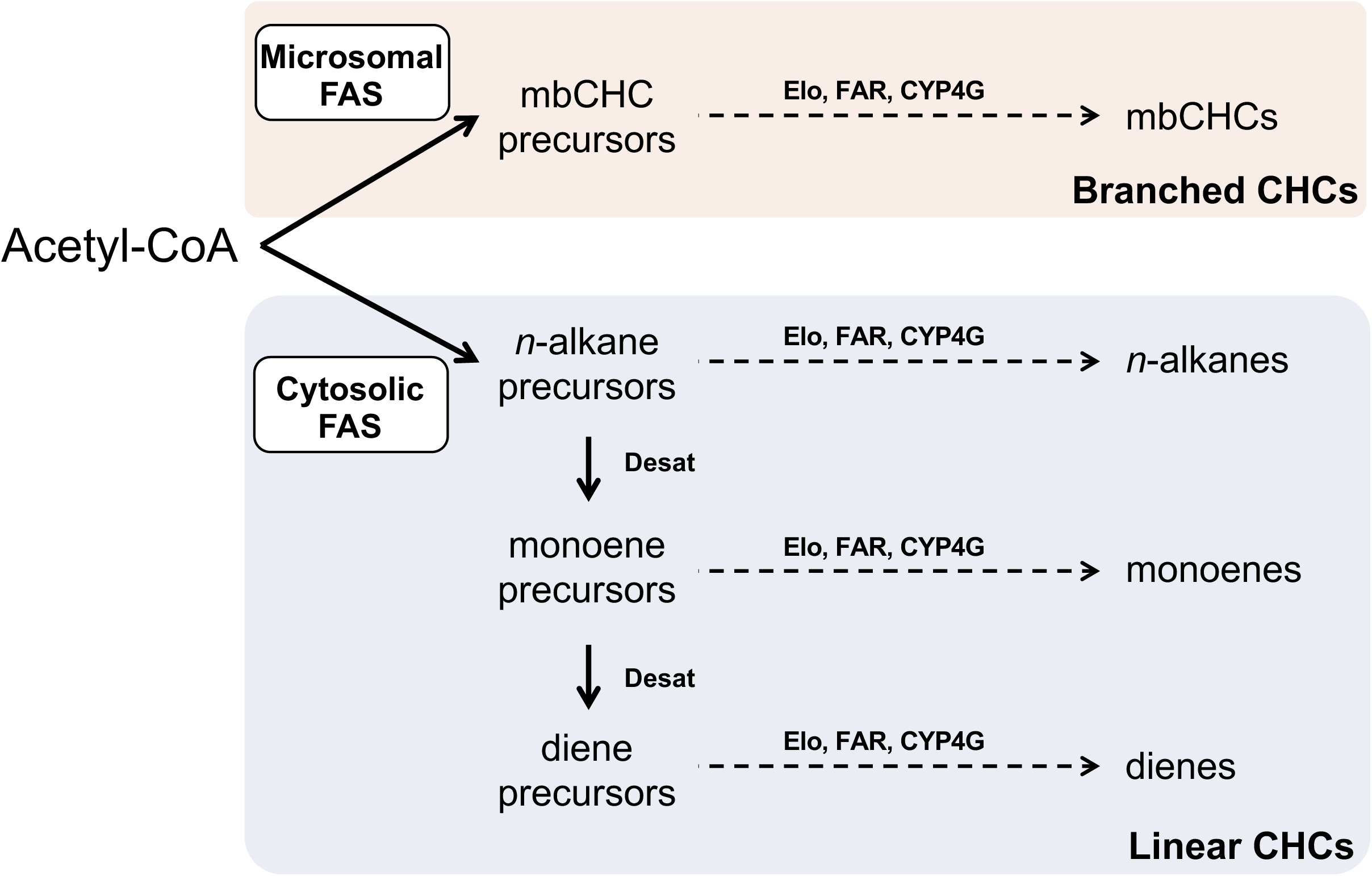
Pathway for the synthesis of branched CHCs (mbCHCs) and linear CHCs (n-alkanes, monoenes, dienes). All CHCs are synthesized via the fatty acyl-CoA synthesis pathway from acetyl-CoA. The pathway splits early due to the action of two fatty acid synthetases (FAS) into the branched CHC pathway or the linear CHC pathway. The precursors of each pathway are modified by synthesis enzymes such as elongases (Elo), reductases (FAR), and the terminal P450 decarbonylase (CYP4G) into CHCs. For linear CHCs, additional enzymes such as desaturases (Desat) incorporate double bonds during synthesis. The diagram was modified from Chung and Carroll, 2015.

**Figure S10.**
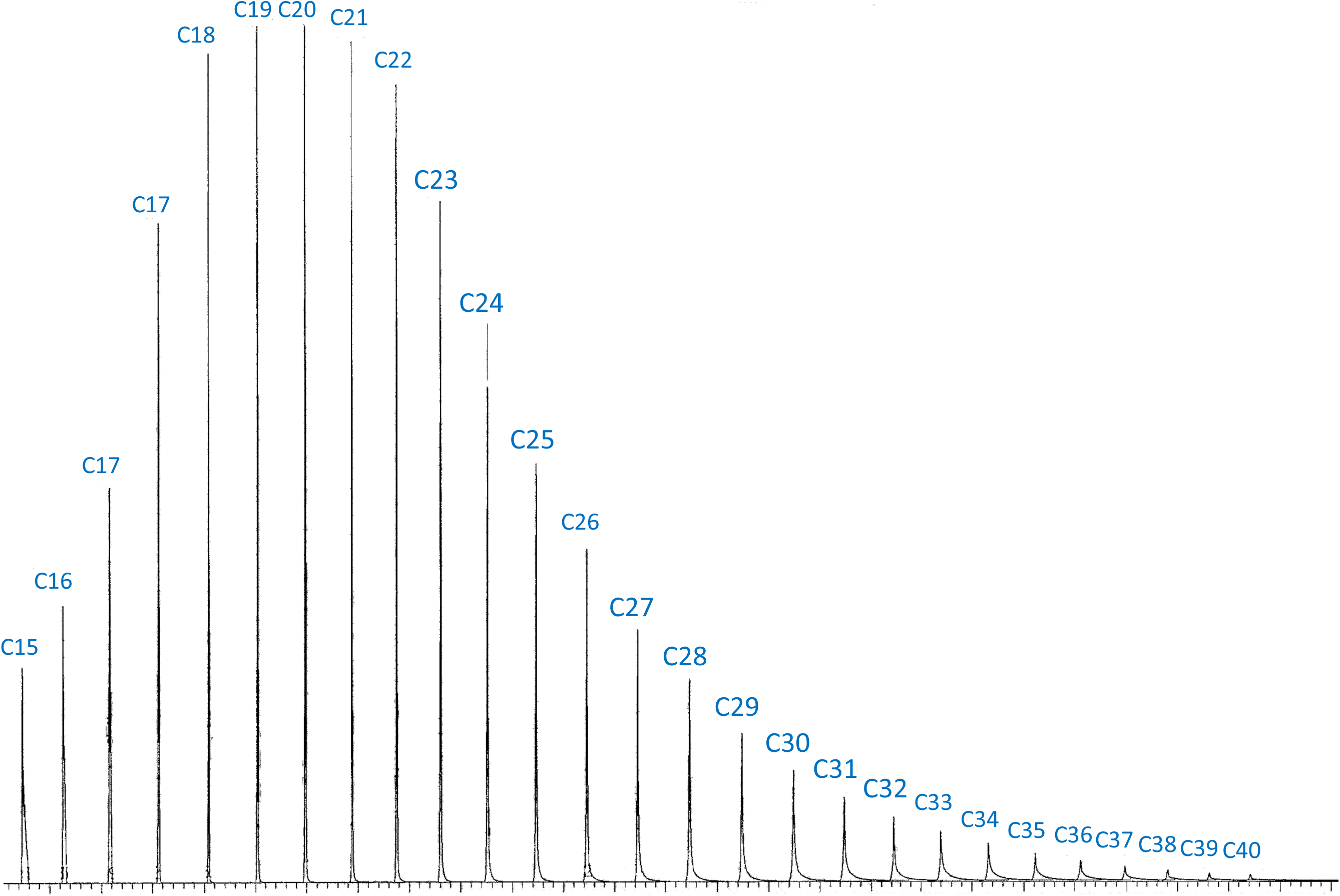
Chromatogram of the authentic standard *n*-alkane mixture which contains 34 *n*-alkanes from C7 to C40 at the same concentration. The authentic standard *n*-alkane mixture (C7-C40; Supelco^®^ 49452-U) was obtained from Sigma-Aldrich (St. Louis, MO). C7 – C14 were excluded in the chromatogram due to the solvent delay in the GC-MS program. The integration of peak areas on longer CHCs is biased to have lower values. To address this issue, we corrected the peak areas of CHCs based on the carbon-chain lengths using the integration from this standard *n*-alkane mixture.

### SUPPLEMENTARY TABLES

**Table S1.** Phylogenetic signals for mbCHCs in *Drosophila* species.

| Sex | 2MeC24 | 2MeC26 | 2MeC28 | 2MeC30 | 2MeC32 | Pagel's $\lambda$ |
| --- | --- | --- | --- | --- | --- | --- |
| Female | 92.2% | 87.9% | 90.9% | 90.4% | 88.8% | 0.75 |
| Male | 93.1% | 90.1% | 91.9% | 91.8% | 92.0% | 0.82 |

**Table S2.** Phylogenetic signals for desiccation resistance in *Drosophila* species.

| Sex | Pagel's $\lambda$ | LogLikelihood ($\lambda$) | Likelihood_Ratio ($\lambda = 0$) | P-value |
| --- | --- | --- | --- | --- |
| Female | 0.91 | -22.4 | 45.2 | < 0.001 |
| Male | 0.97 | -25.4 | 61.7 | < 0.001 |

**Table S3.** Summary of the Phylogenetic Generalized Linear Square (PGLS) models between the longest mbCHCs and desiccation resistance for females and males in 50 *Drosophila* and related species.

| Sex | Term | t value | P value |
| --- | --- | --- | --- |
| Female | Length | 3.2 | 0.002 |
|  | Quantity | -3.6 | < 0.001 |
|  | Interaction | 3.5 | < 0.001 |
| Male | Length | 1.9 | 0.08 |
|  | Quantity | -2.3 | 0.03 |
|  | Interaction | 2.3 | 0.03 |

**Table S4.**
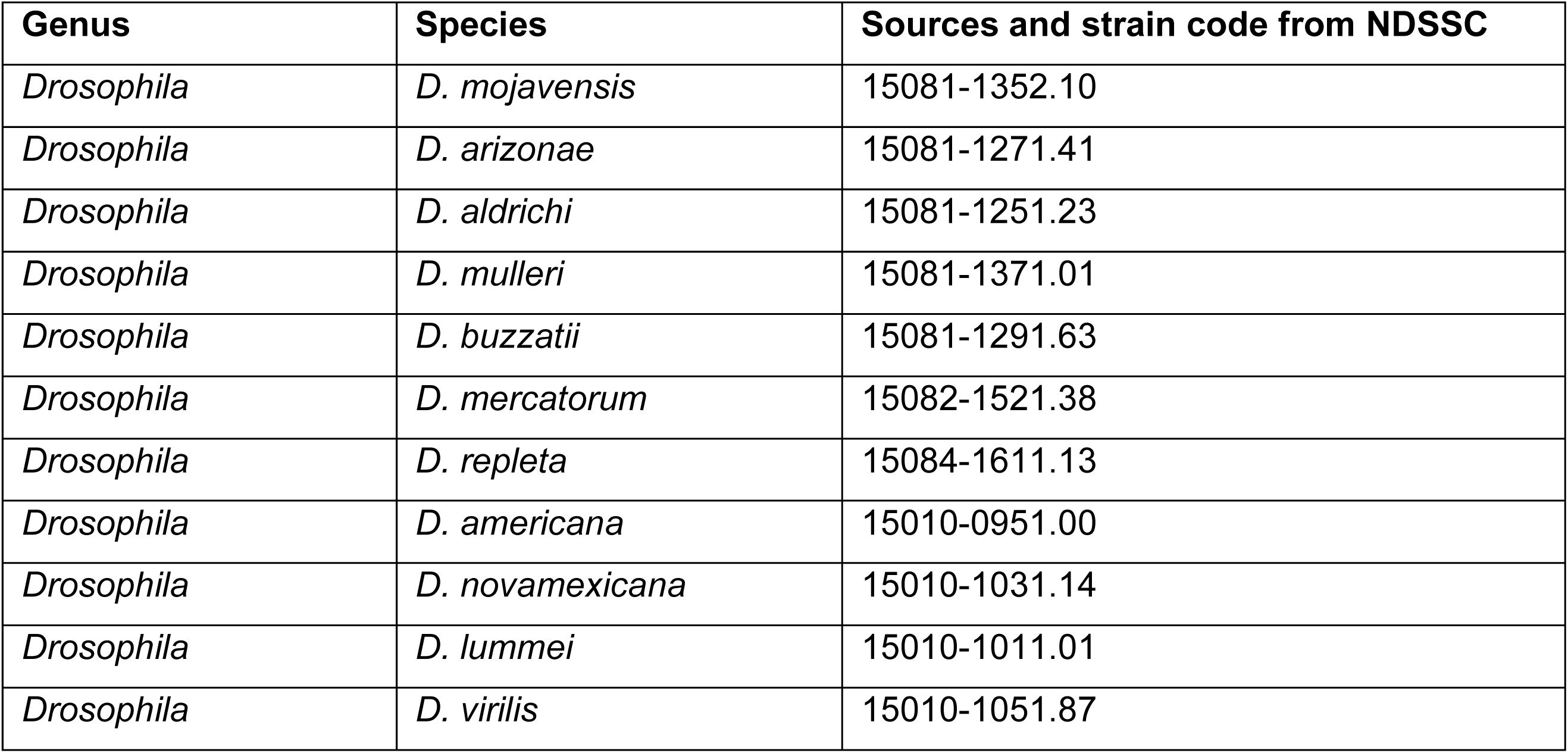
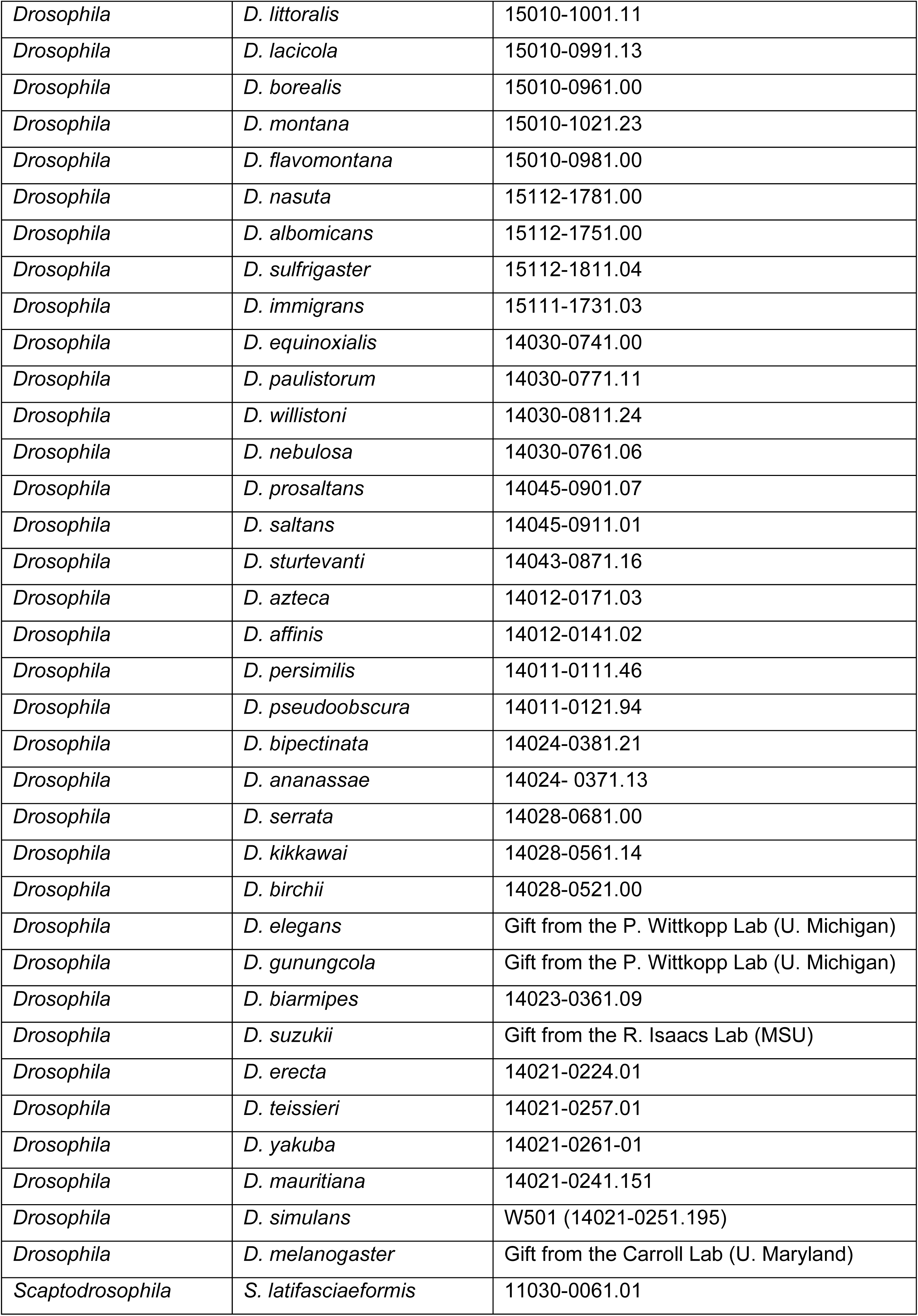
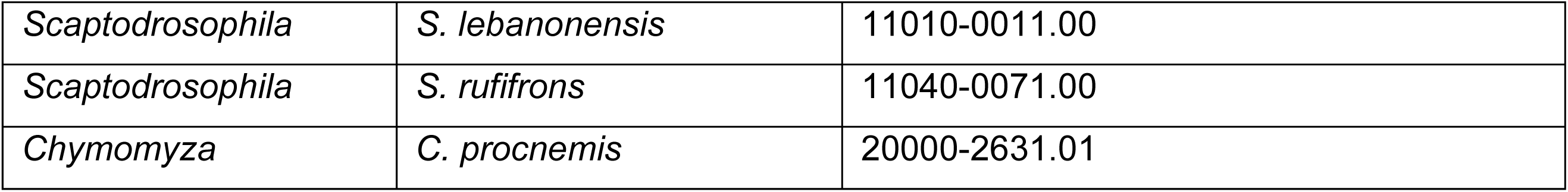
List of species used in this study

## Notes

### Competing Interest Statement

The authors have declared no competing interest.

